# Mutational Landscape of Spontaneous Base Substitutions and Small Indels in Experimental *Caenorhabditis elegans* Populations of Differing Size

**DOI:** 10.1101/529214

**Authors:** Anke Konrad, Meghan J. Brady, Ulfar Bergthorsson, Vaishali Katju

## Abstract

Experimental investigations into the rates and fitness effects of spontaneous mutations are fundamental for our understanding of the evolutionary process. To gain insights into the molecular and fitness consequences of spontaneous mutations, we conducted a mutation accumulation (MA) experiment at varying population sizes in the nematode *Caenorhabditis elegans*, evolving 35 lines in parallel for 409 generations at three population sizes (*N* = 1, 10, 100 individuals). Here, we focus on nuclear SNPs and small indels under minimal influence of selection, as well as their accrual rates in larger populations under greater selection efficacy. The spontaneous rates of base substitutions and small indels are 1.84 × 10^−9^ substitutions and 6.84 × 10^−10^ changes /site/generation, respectively. Small indels exhibit a deletion-bias with deletions exceeding insertions by three-fold. Notably, there was no correlation between the frequency of base substitutions, nonsynonymous substitutions or small indels with population size. These results contrast with our previous analysis of mtDNA mutations and nuclear copy-number changes in these MA lines, and suggest that nuclear base substitutions and small indels are under less stringent purifying selection compared to the former mutational classes. A transition bias was observed in exons as was a near universal base substitution bias towards A/T. Strongly context-dependent base substitutions, where 5’–T and 3’–As increase the frequency of A/T → T/A transversions, especially at the boundaries of A or T homopolymeric runs, manifest as higher mutation rates in (i) introns and intergenic regions relative to exons, (ii) chromosomal cores versus arms and tips, and (iii) germline-expressed genes.

## INTRODUCTION

Spontaneous mutation is central to our understanding of the evolutionary process, given its role as the preeminent source of genetic variation. A detailed understanding of the rate and spectrum of spontaneous mutations is critical for the interpretation of genetic variation in natural populations, the evolutionary dynamics of mutations under the forces of natural selection and genetic drift, the limits to adaptation, the nature of complex human disease and cancer, and the genetic and phenotypic consequences of maintaining populations at small sizes, among others. Because natural variation is the result of an interplay between mutations, genetic drift and natural selection, having a realistic hypothesis for genetic variation in the absence of selection is essential. Furthermore, features of the genome can be shaped by prevailing mutational biases such as base composition, and in turn, the base composition itself can influence mutation rates (Smith et al. 2002; Krasovec et al. 2017). Moreover, mutation rates themselves are not uniformly distributed across genes in the genome. In addition to base composition, variables such as age, replication timing, chromatin organization, and gene expression have been suggested to influence the mutation rate (Hodgkinson and Eyre-Walker 2011).

Mutation accumulation (MA) experiments have a rich history in evolutionary biology since the late 1960s, having provided us a relatively unbiased view of the mutation process by enabling the study of newly originated mutations with minimal interference from the eradicative influence of purifying selection. Replicate lines descended from a single ancestral genotype are evolved independently under extreme bottlenecks each generation to diminish the efficacy of selection, thereby promoting evolutionary divergence due to the accumulation of mutations by random genetic drift. This experimental evolution design of MA experiments circumvents the challenges associated with studying newly arisen mutations in natural or wild populations where strong selection may purge the very mutational variants of interest (reviewed in Halligan and Keightley 2009; Katju and Bergthorsson 2019).

MA experiments typically maintain all replicate lines at the same minimal population size. A variation on this theme, comparing the rate of mutation accumulation between MA lines maintained at different population sizes, enables one to manipulate the strength of selection as a function of population size. In our *C. elegans* MA experiment, all MA lines descended from a single N2 hermaphrodite ancestor, were bottlenecked each generation at *N* = 1, 10, or 100 hermaphrodites (**Supplemental Fig. S1A**) for >400 generations. This experimental design permits a simultaneous investigation of the effects of spontaneous mutation and selection on genetic variation, as well as indirect inferences of the fitness consequences of different classes of mutations. We have previously measured the spontaneous rates and properties of new mutations in the mtDNA genome (Konrad et al. 2017) and nuclear copy-number variants (CNVs) (Konrad et al. 2018) in *C. elegans* under strong genetic drift as well as an increasing efficacy of selection. In both analyses, there was evidence of selection in the larger population size treatments. With regards to the mitochondrial genome, there was no difference in the accumulation of synonymous mutations across different population size treatments, whereas nonsynonymous mutations, frameshifts and deletions accumulated at a higher rate in MA lines maintained at the most extreme population bottleneck of *N* = 1 (Konrad et al. 2017). The accumulation of CNVs in the nuclear genome also showed a significant relationship with population size (Konrad et al. 2018). Gene deletions accumulated at a higher rate in the smallest *N* = 1 populations, and the frequency of gene duplications in the larger populations (*N* =10, 100 individuals) was significantly influenced by gene expression which suggested that (i) high ancestral transcription levels of genes, as well as the (ii) degree of increase in transcript abundance of duplicated genes contribute to the fitness cost of gene duplications.

Here we employ the same set of spontaneous *C. elegans* MA lines comprising three population size treatments (Katju et al. 2015, 2018; Konrad et al. 2017, 2018) and leverage this experimental framework with high-throughput sequencing to identify *de novo* nuclear base substitutions and small indels at a genome-wide scale since the divergence of the MA lines from their common ancestor. With the completion of this study, we are able to (i) offer a comprehensive view of the spontaneous mutation process in *C. elegans*, across both the organellar and nuclear genomes, and all major classes of mutations (base substitutions, small indels and CNVs), (ii) compare our spontaneous mutation rates for nuclear SNPs to previously generated rates that employed older sequencing technologies, (iii) provide one of the first direct, genome-wide estimates of the spontaneous small indel rate for a nematode, and (iv) investigate selective constraints that may impinge on nuclear base substitutions and small indels.

## RESULTS

We sequenced the genomes of 86 *C. elegans* MA lines and their N2 ancestor from a long-term MA experiment with differing population sizes (Katju et al. 2015, 2018; Konrad et al. 2017, 2018). The MA phase of the experiment lasted for 409 generations and comprised three population size treatments wherein a new worm generation was established with *N* = 1, 10 or 100 hermaphrodite worms. 1, 10 or 100 virgin L4 larva(e) were randomly picked to breed in the next generation every four days (**Supplemental Fig. S1A**). For the 20 MA lines (1A–1T) maintained at population size *N* = 1 and the ancestral pre-MA N2 control, the genome of a population of worms derived from one hermaphrodite per line was sequenced (**Supplemental Fig. S1B**). In MA lines comprising larger population sizes, the genomes of four and five individuals were sequenced per *N* = 10 (10 lines; 10A–10J) and *N* = 100 (five lines; 100A–100E) line, respectively. This sequencing design yielded 40 and 25 genomes for the *N* = 10 and *N* = 100 MA lines, respectively (**Supplemental Fig. S1B**). The average read depth was 27.3×, 15.5× and 16.8× per individual genome within the *N* = 1, 10, and 100 population size treatments, respectively. A total of 2,355 single nucleotide polymorphisms (SNPs; **Supplemental Table S1**) and 1,053 small indels (1-100 bp) (**Supplemental Table S2**) were called across all sequenced MA lines (**Supplemental Fig. S2**). Because differing intensities of selection versus drift were hypothesized for the three different population sizes, we analyzed the mutation rates and spectrum separately for each population size treatment.

### Genome-wide estimate of the spontaneous base substitution rate in C. elegans

Single nucleotide substitutions accounted for 1,112 mutations across the *N* = 1 lines, yielding a spontaneous base substitution rate of 1.84 × 10^−9^ /site/generation (**Table 1**; **Supplemental Fig. S2**). The per base substitution rates between the individual *N* = 1 lines range from 1.43 × 10^−9^ to 2.54 × 10^−9^ per generation. The variation among lines was not greater than expected by chance (χ^2^ = 7.8e-10, *df* = 16, *p* = 1) and there was no correlation between mutation rate and the fitness of individual *N* = 1 MA lines (*r* = −0.009, *p* = 0.97). Our estimate of the spontaneous base substitution rate falls within the range previously reported for *C. elegans*, other nematodes and multicellular eukaryotes (**Fig. 1**). However, it is 4.5-fold lower than the earliest direct estimates for *C. elegans* which was based on Sanger sequencing of up to 30 kb of the nuclear genome (Denver et al. 2004). Specifically, our estimate of the nuclear base substitution rate is lower than that reported by Denver et al. (2009) (*t* = 3.76, *p* = 0.004) but higher than that of Denver et al. (2012) (*t* = 3.15, *p* = 0.004) (**Fig. 1**). However, there is no significant difference when the average rate in the N2 strain from the two previous studies (Denver et al. 2009, 2012) is compared to our estimate (*t* = 2.03, *p* = 0.058).

**Table 1.** Summary of the rates of base substitutions and small indels under three population size treatments. Rate estimates for the *N* =1 MA lines represent the spontaneous rate of origin of the various classes of mutations with minimal influence of selection.

| | $N = 1$ | $N = 10$ | $N = 100$ |
| --- | --- | --- | --- |
| $\mu_{bs}$ (/site/generation)<sup>†</sup> | $1.84 \times 10^{-9}$ | $1.95 \times 10^{-9}$ | $1.83 \times 10^{-9}$ |
| $\mu_{indel}$ (/site/generation)<sup>‡</sup> | $6.84 \times 10^{-10}$ | $9.46 \times 10^{-10}$ | $6.95 \times 10^{-10}$ |
| $\mu_{ins}$ (/site/generation)<sup>*</sup> | $1.79 \times 10^{-10}$ | $2.28 \times 10^{-10}$ | $1.90 \times 10^{-10}$ |
| $\mu_{del}$ (/site/generation)<sup>§</sup> | $5.06 \times 10^{-10}$ | $7.18 \times 10^{-10}$ | $5.05 \times 10^{-10}$ |

<sup>†</sup> rate of base substitution
<sup>‡</sup> rate of small indels (insertions and deletions)
<sup>\*</sup> rate of small insertions
<sup>§</sup> rate of small deletions

**Figure 1.**
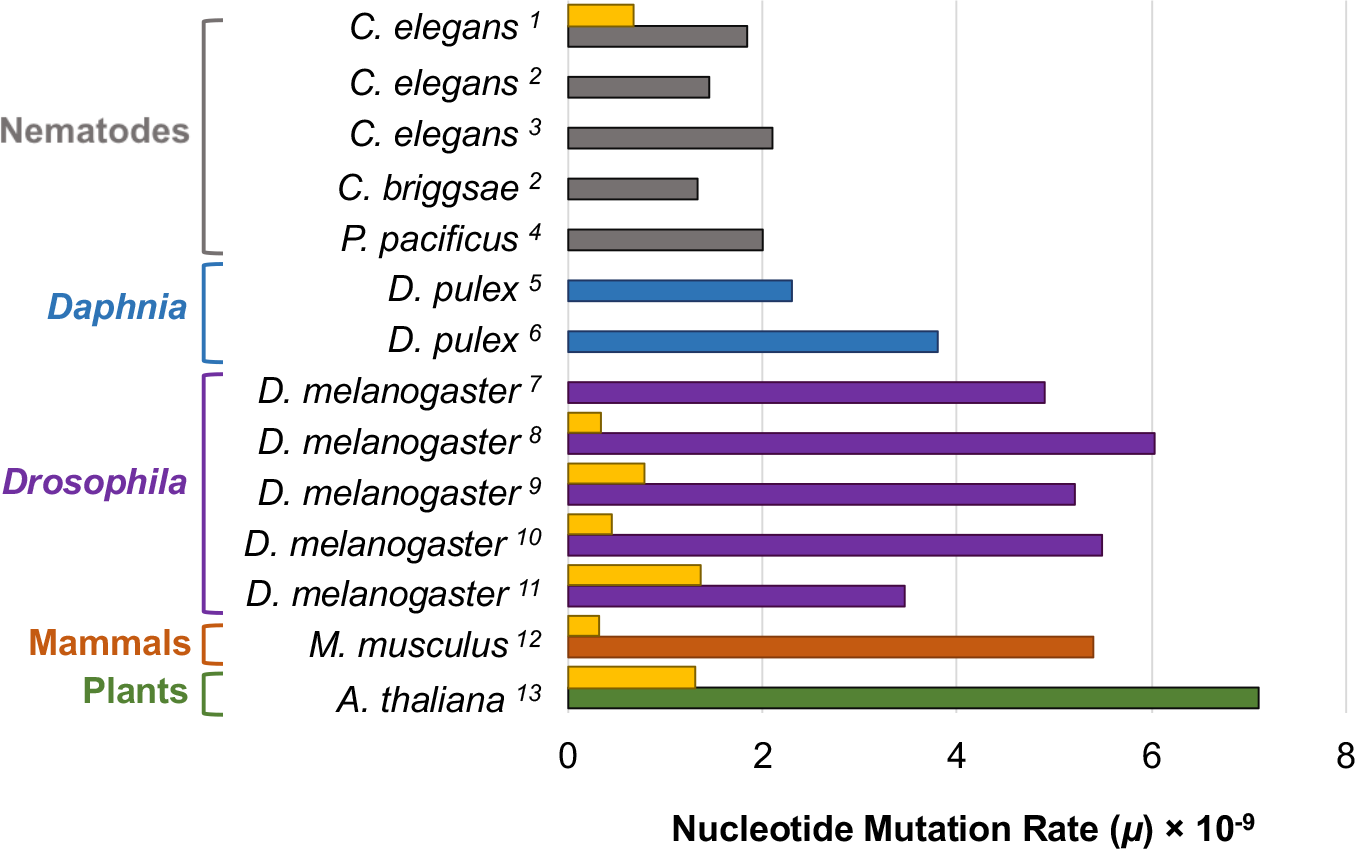
Estimated genome-wide spontaneous base substitution and indel rates for various multicellular eukaryotes. Substitution rates are shown in gray, blue, purple, rust orange and green for nematode, crustacean, insect, mammal, and plant species, respectively. Where available, the yellow bar indicates the indel rate for the corresponding species/study. (Data from: ^1^Current study, ^2^Denver et al. 2012, ^3^Denver et al. 2009, ^4^Weller et al. 2014, ^5^Flynn et al. 2017, ^6^Keith et al. 2016, ^7^Assaf et al. 2018, ^8^Sharp and Agrawal 2016, ^9^Huang et al. 2016, ^10^Schrider et al. 2013, ^11^Keightley et al. 2009, ^12^Uchimura et al. 2015, ^13^Ossowski et al. 2010).

### Estimate of the genome-wide spontaneous indel mutation rate in a nematode and a pronounced deletion-bias

We characterized small insertion and deletion (indel) events as comprising the addition or removal of 100 bp sequences or less, respectively. We detected 357 small indel events in the *N* = 1 lines, resulting in a genome-wide spontaneous indel rate of 6.84 × 10^−10^ /site/generation (**Fig. 1**; **Table 1**; **Supplemental Fig. S2**). Spontaneous indel rates have been reported for *Drosophila melanogaster* (Keightley et al. 2009; Schrider et al. 2013; Huang et al. 2016; Sharp and Agrawal 2016) and *Arabidopsis thaliana* (Ossowski et al. 2010), ranging from 3.38 × 10^−10^ to 1.37 × 10^−9^ /site/generation (**Fig. 1**). Our estimate of the indel rate for *C. elegans* falls within this reported range.

In the *N* =1 MA lines reflecting the spontaneous mutation spectrum, we observed small deletion and insertion rates of 5.06 × 10^−10^ /site/generation and 1.70 × 10^−10^ /site/generation, respectively (**Table 1**). This results in a significant deletion-bias of 2.98 deletions per insertion. This finding is in stark contrast to Denver et al.’s (2004) study that reported a predominance of insertion mutations based on a partial genome analysis (14 – 29 kb) of a different set of *C. elegans N* = 1 MA lines. If all MA lines across our three population size treatments are considered, we observed 519 deletions and 180 insertions resulting in a deletion-bias of 2.88 deletions per insertion. Hence, the deletion-bias is consistent across population sizes (**Supplemental Figs. S3A and S3B**) and deletion rates among all MA lines are significantly higher than insertion rates (**Fig. 2A**; *t* = −9.63, *p* = 3.06 × 10^−12^). The vast majority of indels in our study (67% in *N* = 1 lines) are single-nucleotide insertions or deletions and 76% of the indels are three or fewer nucleotides. The size distribution is also different between insertions and deletions as a greater proportion of deletions relative to insertions exceed two nucleotides (**Fig. 2B**; Wilcoxon test: *W* = 48020, *p* = 5.73 × 10^−7^). This strong deletion-bias, as well as the difference in length distributions between insertions and deletions resulted in a spontaneous net loss of 1,495 bases from the genomes of the *N* = 1 MA lines, an average of 88 bases per genome over the entire experiment, or 0.24 bases per genome per generation.

**Figure 2.**
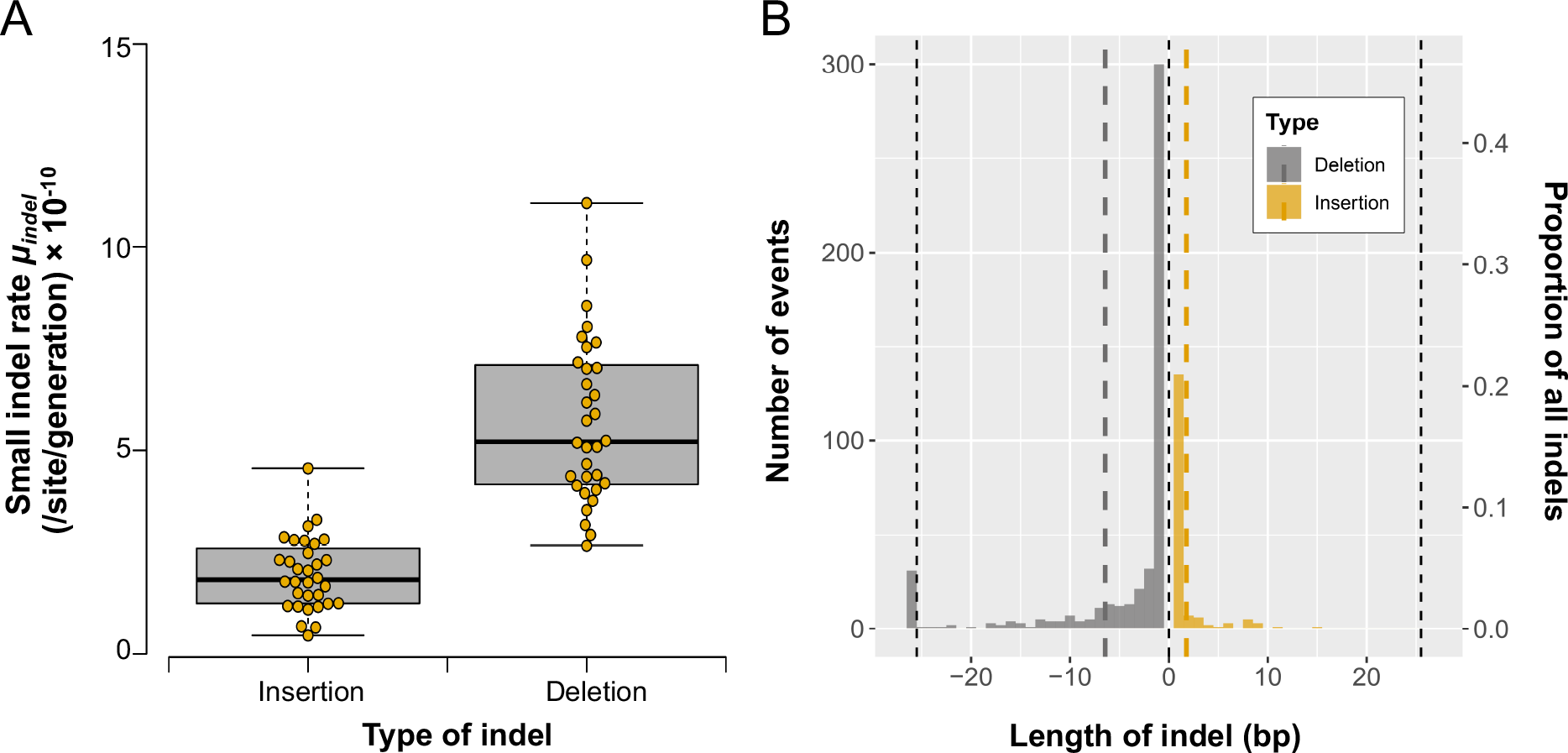
Rates and size distribution of small insertion and deletion events. (A) The deletion rates among all MA lines are significantly higher than insertion rates (*t* = −9.63, *p* = 3.06 × 10^−12^). (B) The size distribution of indels reveals that deletions tend to be larger than insertions (Wilcoxon test: *W* = 48020, *p* = 5.73 × 10^−7^).

### No difference in the base substitution or indel rates between populations of different sizes

Our analysis identified 788 and 455 independent base substitutions in the *N* = 10 and *N* = 100 lines, respectively. The average base substitution rate in the *N* = 10 and *N* = 100 MA lines was 1.95 × 10^−9^ and 1.83 × 10^−9^ /site/generation (**Table 1**), respectively. There is no correlation between population size and the base substitution rate (ANOVA *F* = 0.073, *p* = 0.79; Kendall’s *τ* = 0.0698, *p* = 0.63) (**Fig. 3A**). We identified 227 and 116 independent indel events in the *N* = 10 and *N* = 100 lines, respectively. This yielded average indel rates of 9.46 × 10^−10^ and 6.95 × 10^−10^ /site/generation for the *N* = 10 and *N* = 100 lines, respectively (**Table 1**). As was the case for base substitutions, we found no correlation between population size and the indel rate (ANOVA *F* = 1.17, *p* = 0.29; Kendall’s *τ* = 0.22, *p* = 0.125) (**Fig. 3B**).

**Figure 3.**
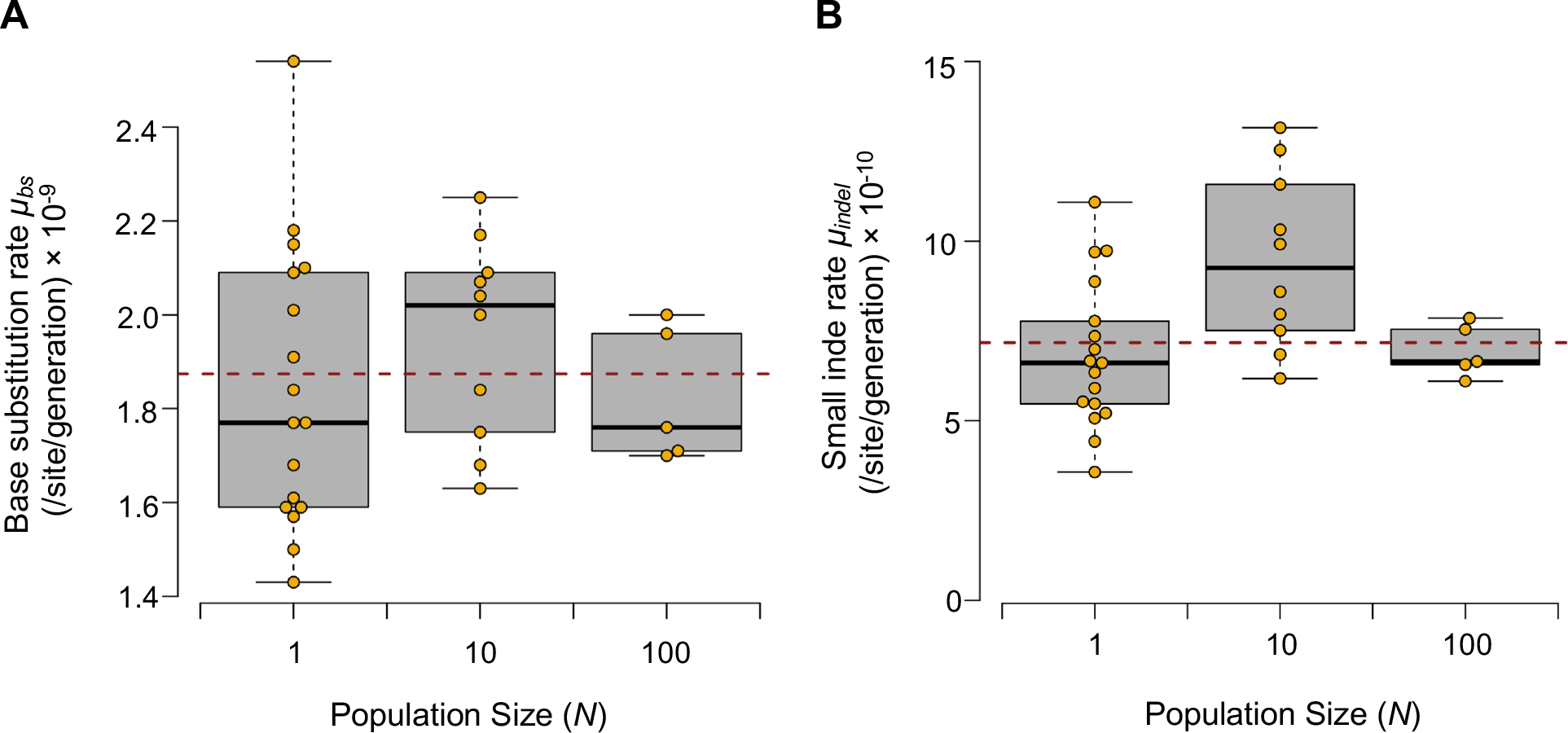
The base substitution and indel rates do not vary with population size. (A) The base substitution rates do not differ significantly between population sizes of *N* = 1, 10, and 100 individuals (ANOVA *F* = 0.073, *p* = 0.79; Kendall’s *τ* = 0.0698, *p* = 0.63). (B) The three population sizes do not differ significantly with respect to the indel rates (ANOVA *F* = 1.17, *p* = 0.29; Kendall’s *τ* = 0.22, *p* = 0.125).

### No discernible difference in the accumulation of nonsynonymous and frame-shift mutations with differing intensity of selection

Natural selection is expected to have greater consequences for the accumulation of nonsynonymous substitutions and frameshift mutations relative to synonymous mutations or mutations in noncoding DNA. Synonymous mutations should be predominantly neutral and we do not expect their rates to vary between different population size treatments. Indeed, there is no difference between the synonymous substitution rates at different population sizes (**Fig. 4A**, ANOVA *F* = 0.04, *p* = 0.84; Kendall’s *τ* = 0.87, *p* = 0.38). In contrast, many nonsynonymous and frameshift mutations are expected to be deleterious and subject to purifying selection in larger populations. However, we did not find significant differences in the nonsynonymous substitution rates (**Fig. 4B**, ANOVA *F* = 0.02, *p* = 0.89; Kendall’s *τ* = 0.27, *p* = 0.79), the combined nonsynonymous substitution and frameshift mutation rates (**Fig. 4C**, ANOVA *F* = 0.07, *p* = 0.79, Kendall’s *τ* = −0.09, *p* = 0.93), or the nonsynonymous/synonymous substitution ratio (*K_a_/K_s_*) between different population sizes (**Fig. 4D**, ANOVA *F* = 1.31, *p* = 0.26, Kendall’s *τ* = −1.1, *p* = 0.27). Furthermore, the median radicality of amino acid changes did not correlate with population size (Kruskal-Wallis *H* = 0.74, *p* = 0.69).

**Figure 4.**
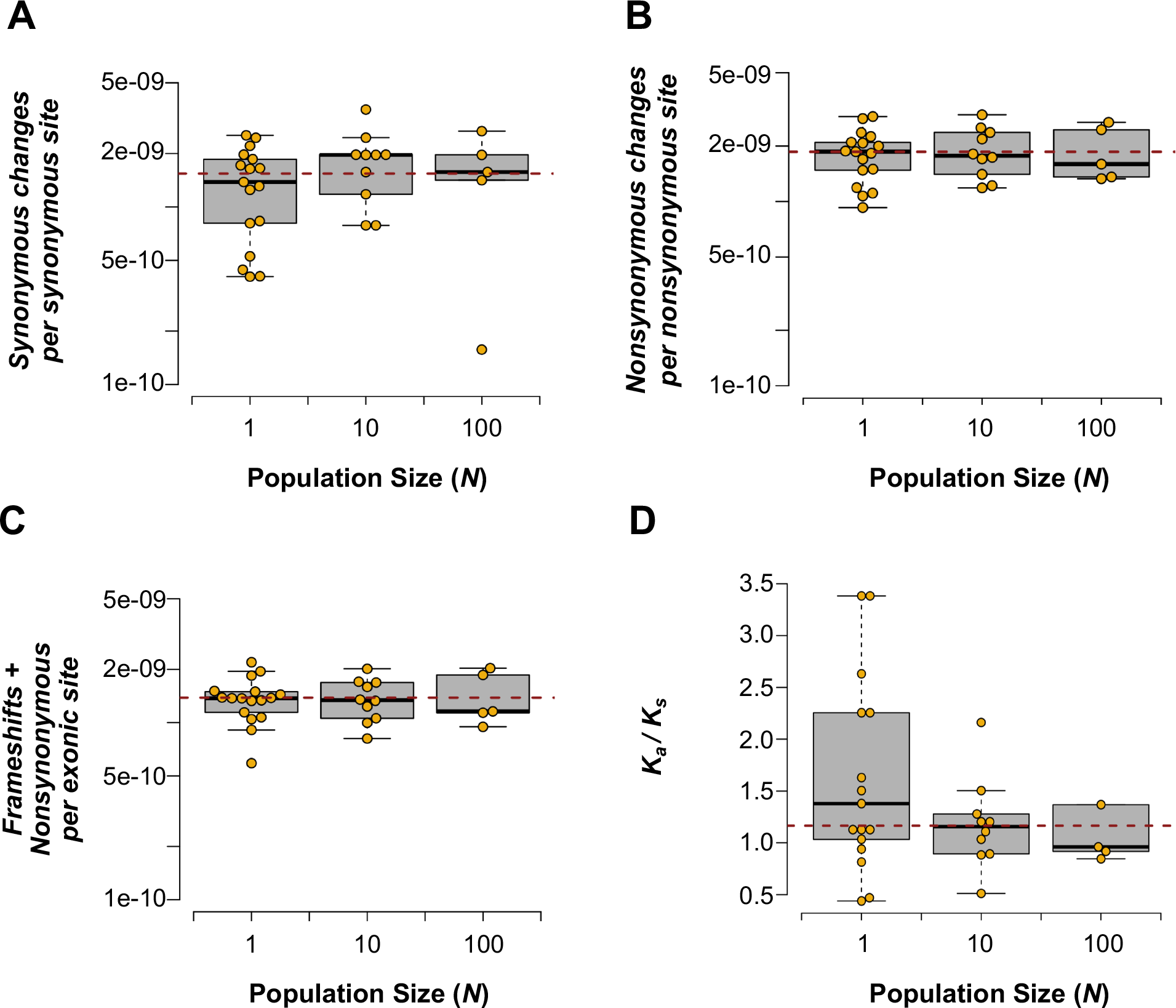
The rates of synonymous and nonsynonymous mutations did not vary with population size. (A) No significant effect of population size is detected in synonymous substitution rates (ANOVA *F* = 0.04, *p* = 0.84; Kendall’s *τ* = 0.87, *p* = 0.38). (B) Nonsynonymous substitution rates do not vary significantly with population size (ANOVA *F* = 0.02, *p* = 0.89; Kendall’s *τ* = 0.27, *p* = 0.79). (C) Pooled nonsynonymous and frameshift mutations rates do not vary significantly with population size (ANOVA *F* = 0.07, *p* = 0.79, Kendall’s *τ* = −0.09, *p* = 0.93). (D) The *K*_*a*_/*K*_*s*_ ratio does not vary with population size (ANOVA *F* = 1.31, *p* = 0.26; Kendall’s *τ* = −1.1, *p* = 0.27).

### Base substitution spectrum exhibits a strong A/T bias

The pattern of base substitutions in the *N* = 1 lines that are under minimal influence of selection should reflect the spontaneous mutation spectrum. The base substitution rate exhibits a strong G/C → A/T mutation bias, primarily driven by G/C → A/T transitions (**Fig. 5**). The mutation rate from a G/C pair to an A/T pair is 2.1, 2.3 and 2.1 × 10^−9^, for the *N* = 1, 10, and 100 lines, respectively. Conversely, the mutation rate from an A/T pair to a G/C pair is 0.56, 0.57 and 0.51 × 10^−9^ for the corresponding population sizes as listed above. Taking *N* = 1 as the best estimate of the mutation rate in the absence of selection, the A/T mutation bias is 3.75. The expected equilibrium G+C-content (GC_eq_), where the number of G/C → A/T mutations equals A/ T → G/C mutations, was calculated as 26% for the *C. elegans* nuclear genome. The *C. elegans* nuclear genome has a G+C-content of 36%.

**Figure 5.**
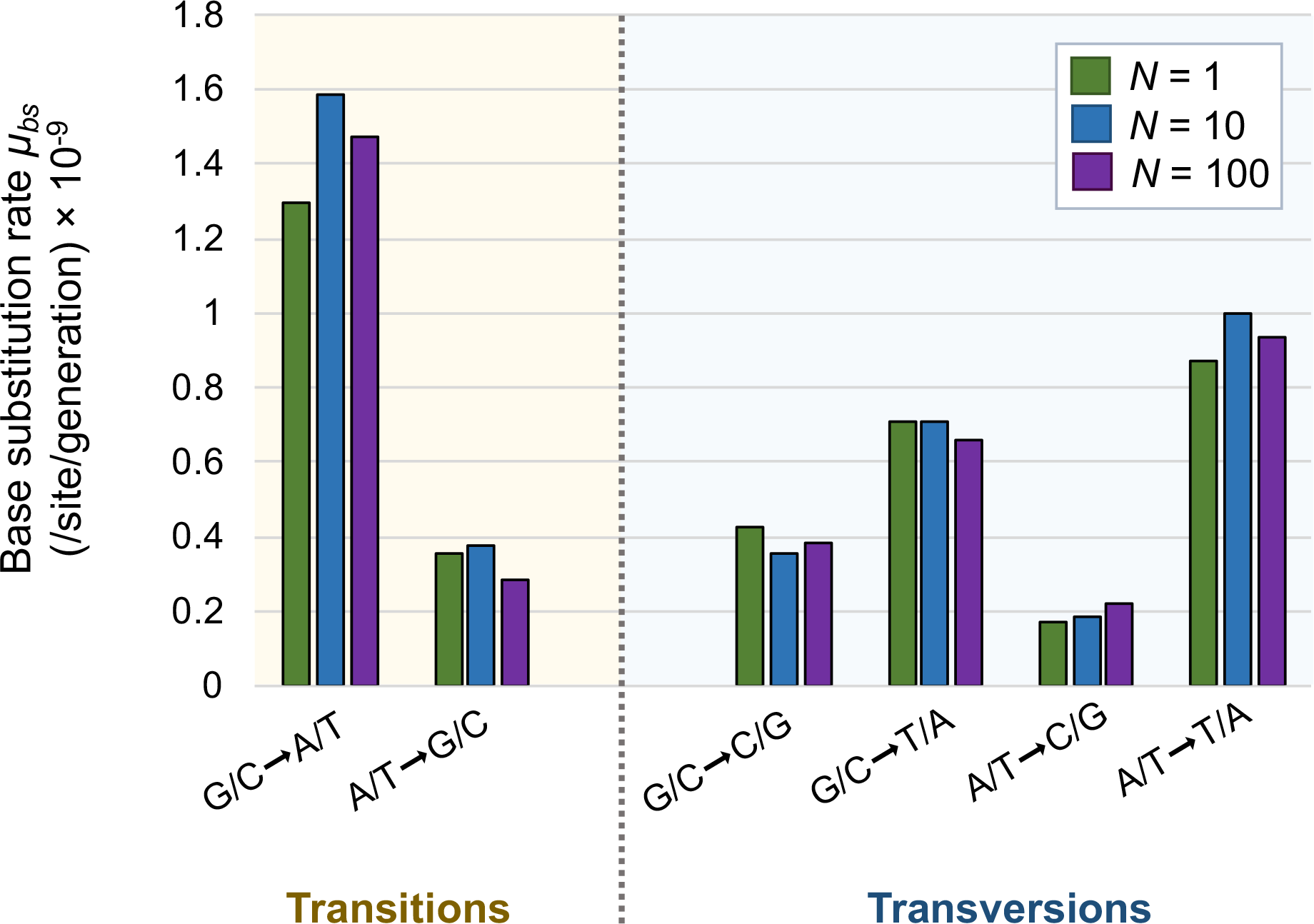
The mutational spectrum at different population sizes. The transition bias is not significantly different from random. The mutational spectrum and the Ts:Tv ratio does not vary with population size (*F* = 0.016, *p* = 0.9).

Base substitutions in the *N* = 1 lines exhibit a slight but nonsignificant transition bias, leading to a transition-transversion ratio (Ts:Tv) of 0.64 (*N* = 1 line specific values range from 0.36–1.04). If all mutations between the four nucleotides are equally likely, the expected transition bias is 0.5. The relative overrepresentation of transitions compared to transversions is therefore 0.64/0.5, or 1.28. The relative overrepresentation of transitions in the *N* = 10 and 100 lines is 1.41 and 1.28, respectively, and the Ts:Tv ratio does not vary with population size (*F* = 0.016, *p* = 0.9). The lack of a strong transition bias is partly due to high rates of A/T → T/A transversions in introns and intergenic regions. If we analyze the transition bias in coding and noncoding sequences separately, the relative overrepresentation of transitions is 1.93 in exons and 1.14 in introns in the *N* = 1 lines.

### Strong context-dependence of A/T → T/A tranversions in noncoding DNA

Compared to previous studies, our data indicate a greater frequency of A/T → T/A transversions. The majority of these mutations are flanked by A and T base pairs on each side and occur more frequently in introns and intergenic regions compared to exons (**Fig. 6A**). A/T → T/A transversions are particularly common in introns and intergenic regions when the focal nucleotide is flanked by a 5’–T and a 3’–A. A flanking 5’–A and 3’–T also appears to elevate the rate of A/T → T/A transversion (**Fig. 6A**). Additionally, these substitutions primarily occur on the boundaries of homopolymeric runs of seven to 11 bases of either adenines or thymines (**Fig. 6B**).

**Figure 6.**
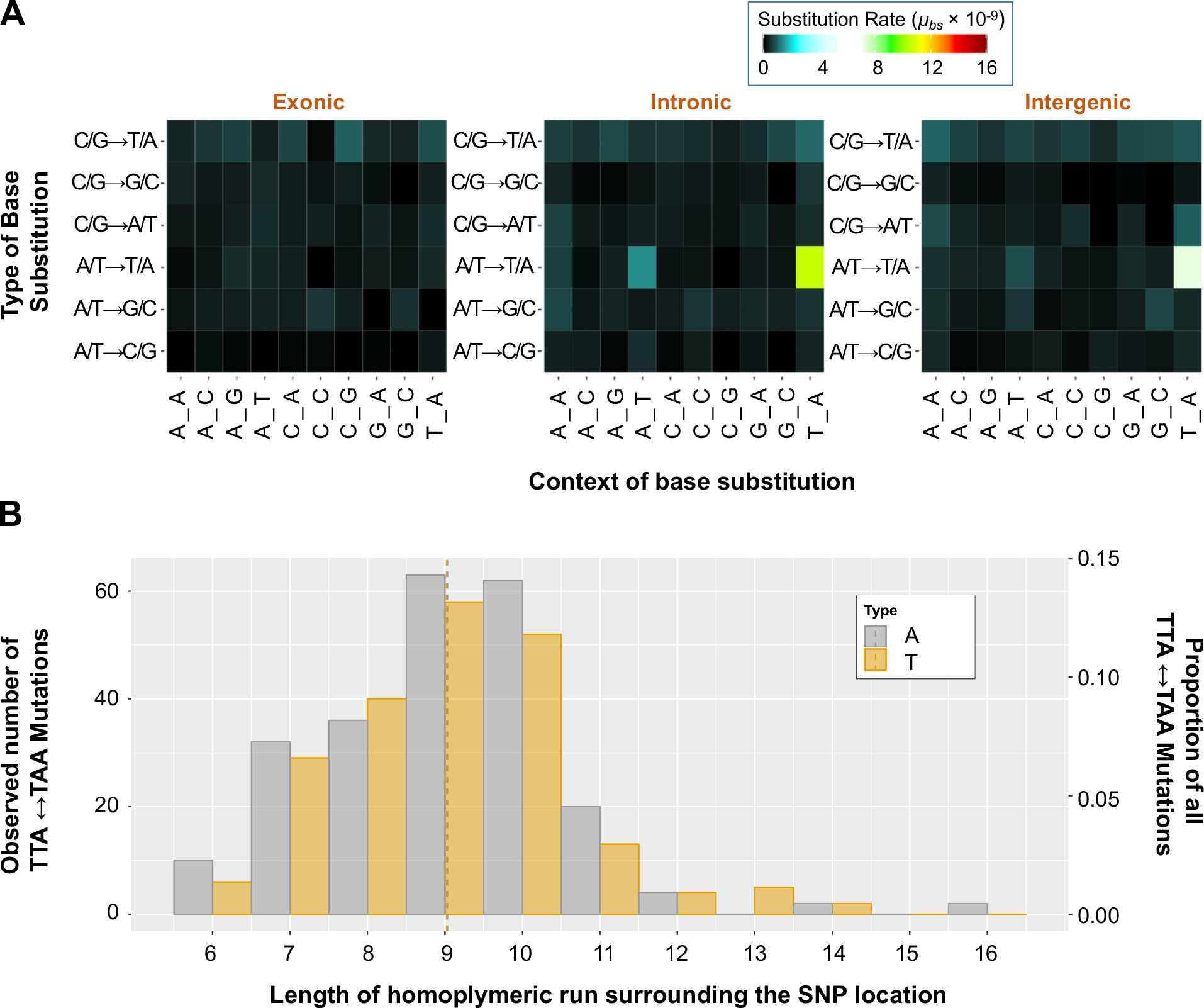
Context-dependence of base substitutions. (A) The vast majority of mutations in intron and intergenic are regions are 5’–TTA–3’ ↔ 5’–TAA–3’ transversions. (B) Substitutions occurring at boundaries of A or T homopolymeric runs are responsible for the disproportionate contribution of A/T→ T/A transversions. The A→ T and T→A transversions are equally frequent in homopolymeric runs, consistent with the absence of a strand bias.

### Elevated base substitution rate in chromosomal arms relative to cores

There was no significant effect of population size on the base substitution rate either at the interchromosomal or intrachromosomal level. Hence, much of the subsequent analysis of the distribution of base substitutions across the *C. elegans* genome will be based on the pooled results from all of the MA lines (*N* = 1, 10, and 100 populations). The nucleotide substitution rates were analyzed in a three-way ANOVA for chromosomes (five autosomes, and one sex chromosome), functional regions (exons, introns and intergenic regions) and recombination zones (arms, cores and tips). The nucleotide substitution rates did not vary significantly between chromosomes (**Fig. 7A**, *F* = 0.86, *p* = 0.51). There is a significant difference between the nucleotide substitution rates in exons, introns and intergenic regions (**Fig. 7B**, *F* = 6.51, *p* = 0.0015). The substitution rate in introns is significantly higher than that in exons (2.25 × 10^−9^ /site/generation, and 1.51 × 10^−9^ /site/generation, respectively; Tukey’s multiple comparisons of means, *p* = 0.001), whereas the nucleotide substitution rates in intergenic regions (1.82 × 10^−9^ substitutions/site/generation) falls between that of introns and exons and is not statistically different from either one. The chromosomal arms comprise 46% of the *C. elegans* genome and are marked by a higher incidence of repetitive elements, lower gene densities, and increased recombination. Chromosomal cores comprising 47% of the genome have higher gene densities, lower repetitive element content, and lower recombination rates. Chromosomal tips are much shorter sections at the ends of chromosomes (7% of the genome) which are not thought to experience recombination (Barnes et al. 1995; Rockman and Kruglyak 2009). The per nucleotide substitution rates differ significantly between chromosomal arms, cores, and tips (**Fig. 7C**; *F* = 6.62, *p* = 0.0014). The nucleotide substitution rate is higher in arms than cores (2.18 × 10^−9^ /site/generation, and 1.58 × 10^−9^ /site /generation, respectively; Tukey’s multiple comparisons of means, *p* = 0.0019), but arms and tips (2.18 × 10^−9^ /site/generation, and 1.96 × 10^−9^ /site/generation, respectively) do not differ significantly in their substitution rates (Tukey’s multiple comparisons of means, *p* = 0.82). The difference in base substitution rates between the arms and the cores is evident for coding and noncoding sequences alike (**Figure 7D**).

**Figure 7.**
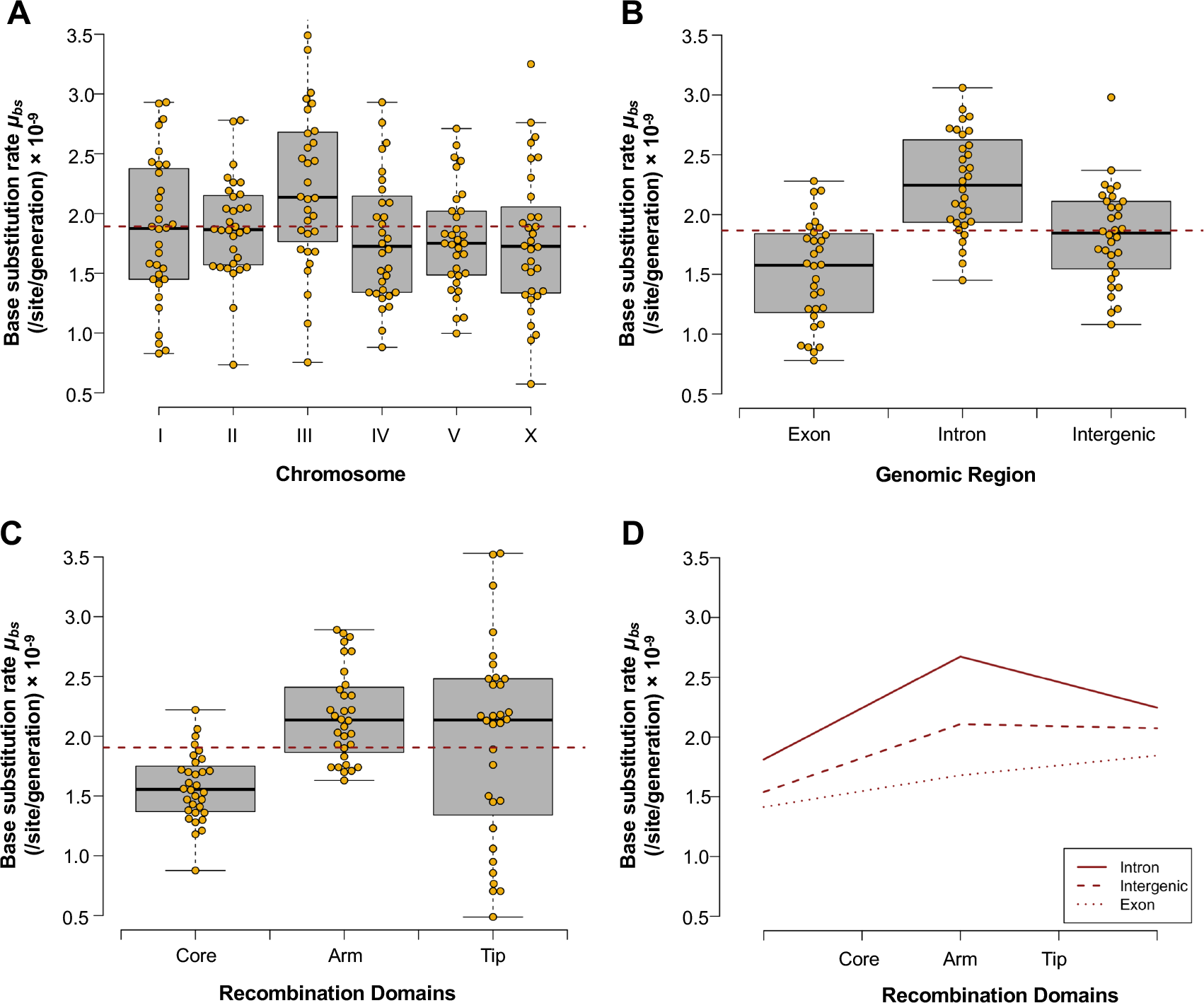
Variation in base substitution rates across different genomic regions. (A) There was no significant difference in the base substitution rate between chromosomes (*F* = 0.86, *p* = 0.51). (B) The base substitution rates differ significantly between exons, introns, and intergenic regions (*F* = 6.51, *p* = 0.0015). (C) Base substitution rates are significantly different between chromosomal arms, cores, and tips (*F* = 6.622, *p* = 0.0014). (D) A lower base substitution rate in cores relative to arms and tips applies to exons, introns and intergenic regions.

### A/T and G/C homopolymeric runs differ in their mutational properties

The number of single nucleotide A or T indels are as expected in the absence of strand bias (**Fig. 8A**). Similarly, G or C single nucleotide indels do not show any evidence of strand bias and occur in roughly equal frequency (**Fig. 8A**; Fisher’s Exact: *p* = 0.508). Furthermore, there is no difference in the spectrum of indels between different population sizes (**Fig. 8B**).

**Figure 8.**
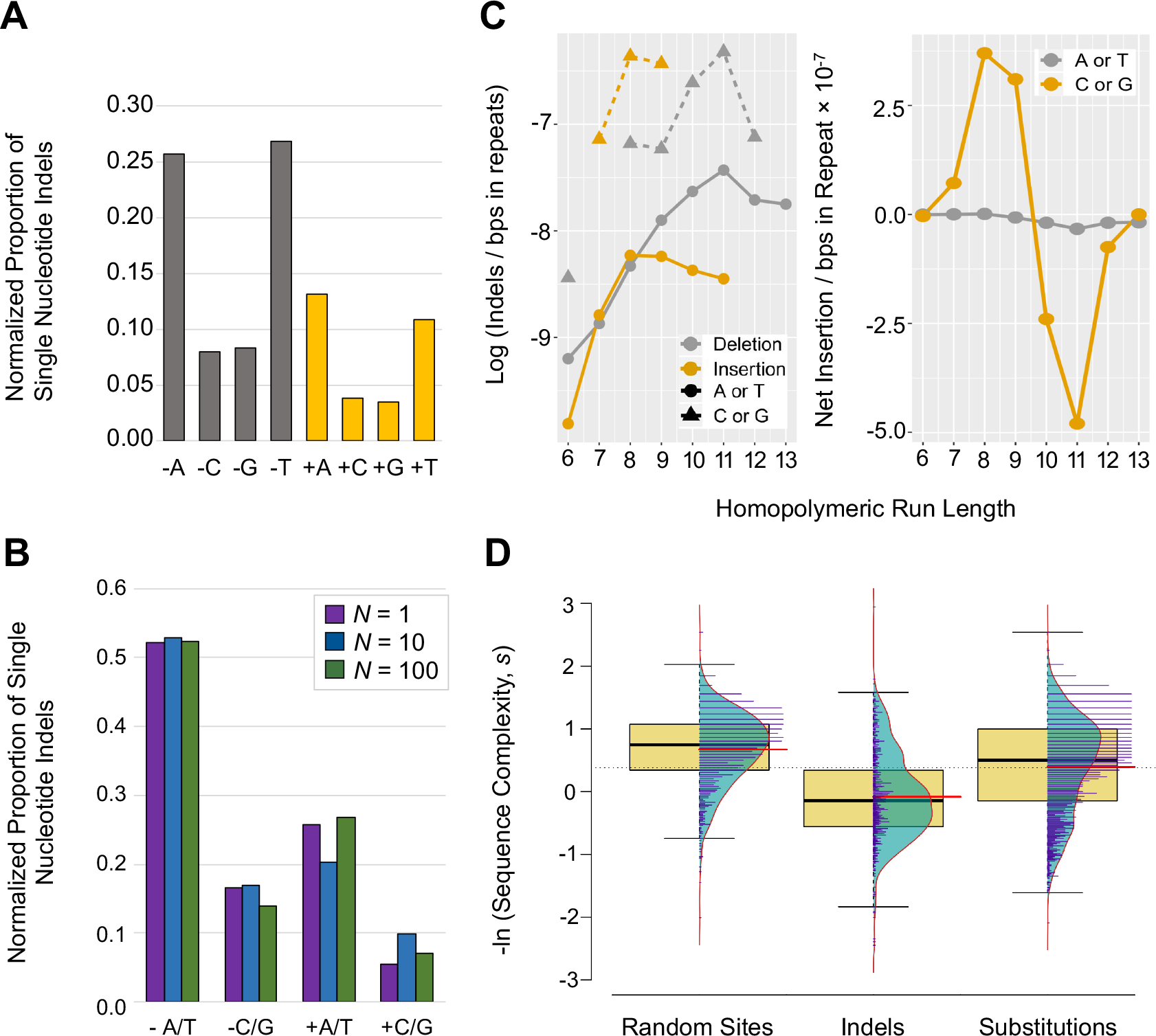
Different rates and patterns of A/T and G/C indels in homopolymeric runs. (A) The number of single nucleotide A or T indels are almost identical and G or C indels are also equally frequent as expected in the absence of strand bias in the indel calls. (B) There is no difference in the frequency of different kinds of single nucleotide indels between different population sizes. (C) G/C homopolymeric runs have higher indel rates than A/T homopolymeric runs. The frequency of A/T indels rises with increasing length of a homoplymeric but then tapers off. The deletion-bias is more pronounced for A/T indels in longer runs as the deletion rates tend to be higher than the insertion rates in long A/T homopolymeric runs. Longer runs of G/C have higher deletion rates than short G/C runs whereas shorter G/C runs have increased insertion rates relative to long runs. (D) The mean sequence complexity surrounding indels is significantly lower than for both random sites in the genome (*t*-test: *t* = −17.03, *p* < 2.2 × 10^−16^), and sequence surrounding base substitutions (*t*-test: *t* = −10.28, *p* < 2.2 × 10^−16^).

While A/T indels are more common across the genome, the G/C indel rates are higher than A/T indel rates after standardizing the rates by mutational opportunity (**Fig. 8C**). The rates of indels in runs of As and Ts increases with the length of the run (**Fig. 8C**). Deletion rates tend to be higher than insertion rates in long A/T homopolymeric runs, and they show similar tendencies as a function of the length of a run. Similarly, longer runs of G/C have higher deletion rates than short G/C runs. In contrast, shorter G+C runs have increased insertion rates relative to long runs (**Figure 8C**). The mean complexity of the sequence that incurred indels is significantly lower than both (i) random sites in the genome (*t*-test: *t* = −17.03, *p* < 2.2 × 10^−16^) and (ii) sequences that incurred nucleotide substitution (*t*-test: *t* = −10.28, *p* < 2.2 × 10^−16^). This is likely due to the propensity of indels to occur mainly in A+T-rich regions, which are by nature of low complexity (**Fig. 8D**).

### Intrachromosomal location significantly affects the small indel rate

The effect of chromosomal location on the indel rates mirrors that of base substitutions. There were no significant interactions between the effects of the chromosome and chromosomal region (three-way ANOVA: *F* = 1.36, *p* = 0.19), the chromosome and the coding content (three-way ANOVA: *F* = 0.94, *p* = 0.5), the chromosomal region and coding content (three-way ANOVA: *F* = 0.48, *p* = 0.75), or all three (three-way ANOVA: *F* = 0.78, *p* = 0.74). The indel rates are not significantly different between individual chromosomes (**Fig. 9A**; Kruskal-Wallis: *H* = 9.01, *p* = 0.11; three-way ANOVA: *F* = 2.13, *p* = 0.06). As was the case for base substitutions, the indel rates differ significantly between exons, introns, and intergenic regions (**Fig. 9B**; three-way ANOVA: *F* = 20.07, *p* = 2.45 × 10^−9^; Kruskal-Wallis: *H* = 50.20, *p* = 1.26 × 10^−11^). Indel rates were observed to be the lowest for exonic regions. Intronic and intergenic regions had higher indel rates, likely attributable to these regions containing different amounts of low complexity sequence. Furthermore, the indel rates differ between chromosomal arms, cores, and tips (**Fig. 9C**; three-way ANOVA: *F* = 3.74, *p* = 0.24; Kruskal-Wallis: *H* = 18.79, *p* = 8.33 × 10^−5^). While no significant indel rate differences are detected between arms and tips (*t*-test: *t* = 0.71, = 0.48; Mann-Whitney: *U* = 545, *p* = 0.67), indel rates are significantly lower in cores than in chromosomal arms (*t*-test: *t* = 5.169, *p* = 3.51 × 10^−6^; Mann-Whitney: *U* = 854, *p* = 4.53 × 10^−6^). The low indel rates in the cores compared to the arms and tips were detected for all functional regions (exons, introns and intergenic regions) (**Fig. 9D).**The distribution of indels across the chromosomal regions does not differ significantly between population size treatments (**Supplemental Fig. S4**; Fisher’s Exact Test: *p* = 0.74).

**Figure 9.**
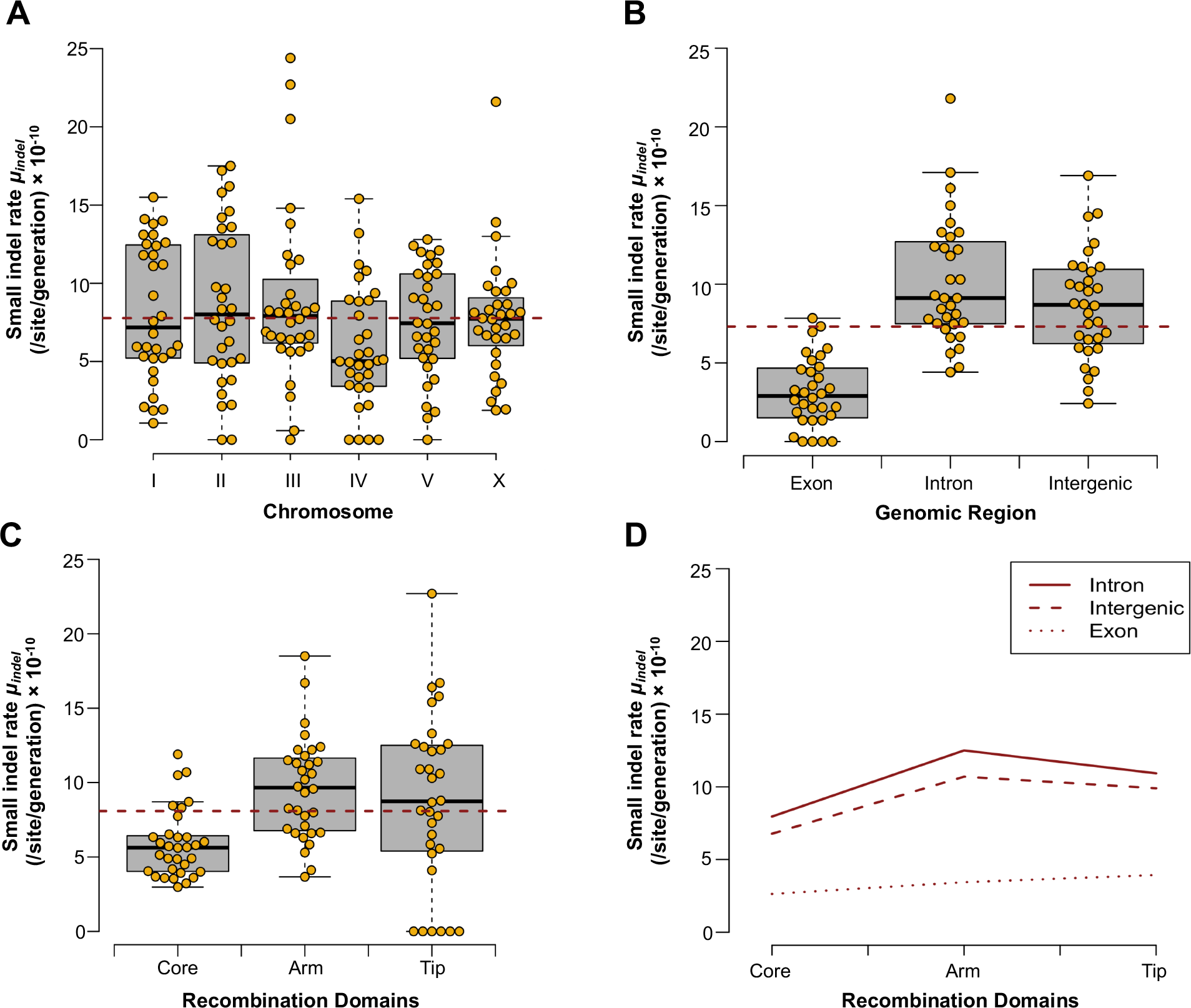
Variation in small indel rates across different genomic regions. (A) There was no significant difference in the small indel rate between chromosomes (Kruskal-Wallis: *H* = 9.01, *p* = 0.11; three-way ANOVA: *F* = 2.13, *p* = 0.06). (B) The indel rate differs significantly between exons, introns, and intergenic regions (Kruskal-Wallis: *H* = 50.2, *p* = 1.26 × 10^−11^, three-way ANOVA: *F* = 20.07, *p* = 2.45 × 10^−9^). (C) The indel rates are significantly different between chromosomal arms, cores, and tips (Kruskal-Wallis: *H* = 18.79, *p* = 8.33 × 10^−5^, three-way ANOVA: *F* = 3.74, *p* = 0.24). (D) A lower indel rate in cores compared to arms and tips applies to exons, introns and intergenic regions.

### Germline expressed genes have higher mutation rates than non-germline expressed genes

The transcription of a gene has the potential to influence its mutation rate and some studies have found a positive association between transcription and mutation rate (Hudson et al. 2003; Alexander et al. 2012; Kim and Jinks-Robertson 2012). In order to determine whether germline expression of *C. elegans* genes is correlated with the mutation rate, we classified the protein-coding genes into germline expressed and non-germline expressed genes using published results (Wang et al. 2009). The substitution rate across all MA lines is significantly higher in germline expressed genes than in non-germline expressed genes (**Fig. 10A**; two-way ANOVA: *F* = 12.05, *p* = 0.0007). Chromosomal cores are more gene-rich than chromosomal arms, and we previously detected a significant difference in substitution rates between those two regions. Moreover, there is a significant interaction between germline expression and the recombination domain (**Fig. 10B**; two-way ANOVA: *F* = 12.8, *p* = 0.0007). With respect to the core regions, there was no difference in the mutation rates of germline and non-germline expressed genes. In contrast, germline expressed genes have higher mutation rates than non-germline genes when residing in the chromosomal arms and tips.

**Figure 10.**
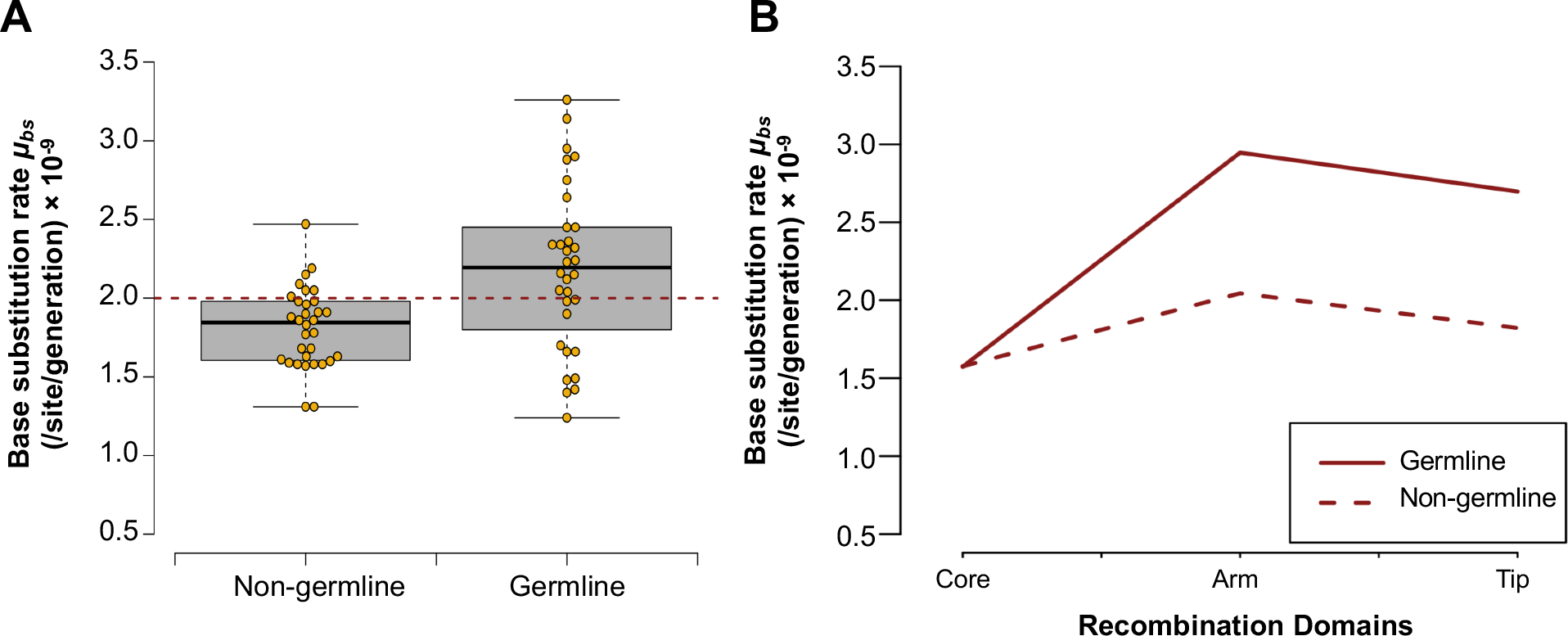
Germline expressed genes have higher mutation rates than non-germline expressed genes. (A) Base substitution rate distributions differ significantly between genes with germline versus non-germline expression (*F* = 12.05, *p* = 0.0007). (B) Germline expressed genes located in chromosomal arms and tips have higher mutation rates than non-germline genes in the same recombination domain (*F* = 12.8, *p* = 0.0007).

### Context-dependent A/T → T/A transversions contribute to intrachromosomal variation in substitution rates

There are significant differences in the frequency of homopolymeric runs between coding and non-coding DNA. Because strongly context-dependent A/T → T/A transversions occur frequently at the boundaries of A/T homopolymers, we tested if any of the positional or transcription related differences in mutation rate could be accounted for by these A/T → T/A transversions. If all A/T → T/A transversions are excluded from the analysis, we no longer observe significant differences in mutation rates between (i) exons and non-coding DNA (**Fig. 11A**), nor (ii) between germline and non-germline transcribed genes (**Fig. 11B**). In contrast, there still exists significant mutation rate variation among chromosomal cores, arms and tips despite the exclusion of A/T → T/A transversions (**Fig. 11C**; ANOVA *F* = 3.9, *p* = 0.024). This variation is primarily due to a significant difference in mutation rates between chromosomal cores and arms (Tukey’s multiple comparisons of means, *p* = 0.02). In sum, the nonrandom distribution of mutable motifs can account for the differences between coding and non-coding DNA, as well as transcription-related differences in mutation rates, and they contribute to the differences in mutation rates between cores, arms and tips. However, the difference in mutation rate between cores, arms and tips are not fully explained by context dependent A/T → T/A transversions. Thus, the higher rates of mutations in arms compared with cores could also be due to higher recombination frequency.

**Figure 11.**
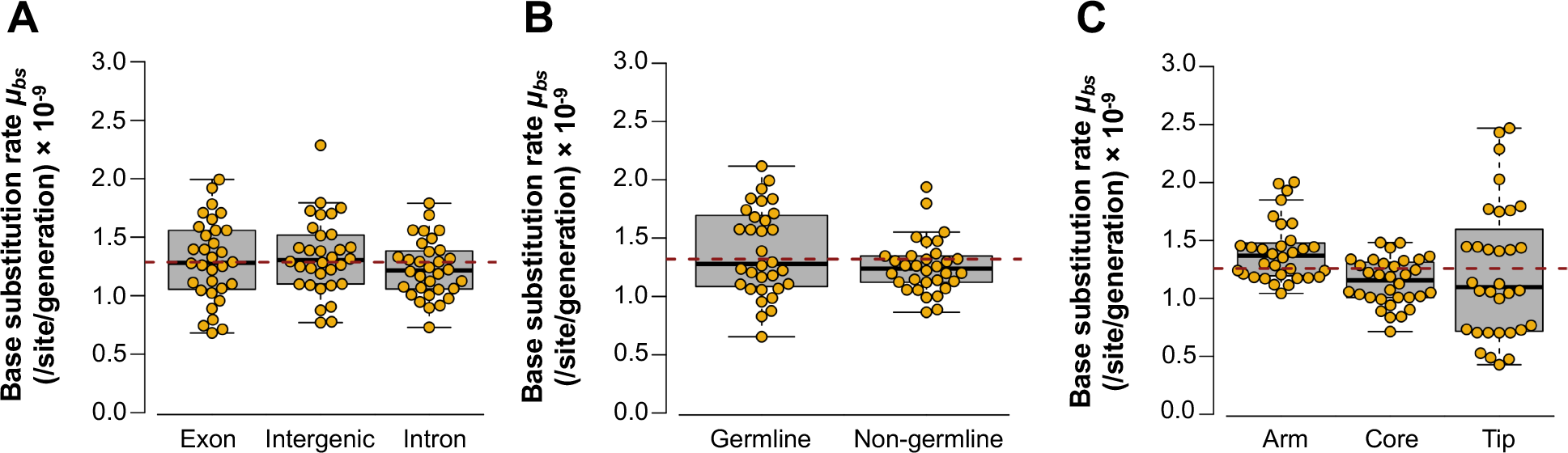
Comparison of mutation rates with respect to genome position and germline transcription when A/T to T/A transversions are excluded from the data. (A) No difference in base substitution rates among exons, introns and intergenic regions (ANOVA *F* = 0.91, *p* = 0.41). (B) No difference in base substitution rates between germline and non-germline expressed genes (*t* = 1.6, *p* = 0.12; Kendall’s *τ* = 0.27, *p* = 0.79). (C) Significant variation in base substitution rates among chromosomal cores, arms and tips (ANOVA *F* = 3.878, *p* = 0.024).

## DISCUSSION

MA experiments typically consist of passaging experimental replicate lines through a minimum population bottleneck in each generation of the experiment. Contrastingly, our *C. elegans* MA experiment comprised three population size treatments aimed at assessing the rates of origin of diverse classes of mutations and their differential accumulation under varying regimes of natural selection. We have previously assessed the phenotypic consequences of mutation and selection under benign laboratory (Katju et al. 2015) and osmotic stress conditions (Katju et al. 2018). In addition, we have employed modern genomic approaches to investigate the interplay of mutation and selection on mtDNA SNPs and small indels (Konrad et al. 2017) as well as nuclear copy-number variants (Konrad et al. 2018). In this study, we additionally investigated two additional major classes of mutational variants in the nuclear genome, namely SNPs and small indels to provide a comprehensive picture of the spontaneous mutation process in *C. elegans* through the lens of experimental evolution.

The *N* = 1 lines provide the baseline for the spontaneous rate of origin of different classes of mutations and the expected rate of neutral evolution. In this study, the spontaneous rate of origin of nuclear base substitutions (*μ*_*bs*_) and small indels of <100 bp length (*μ*_*indel*_) in *C. elegans* were determined to be 1.84 × 10^−9^ substitutions/site/generation and 0.68 × 10^−9^ indels/site/generation, respectively. Hence, the rate of accumulation of nuclear SNPs exceeds that of small nuclear indels by approximately three-fold. Based on this study and our preceding mtDNA genome analysis on the same set of MA lines (Konrad et al. 2017), we find that the spontaneous rates of different classes of mutations per nucleotide in *C. elegans* range from 10^−10^ to 10^−8^ per base per generation, representing a ∼90-fold difference. This relationship can be expressed as follows: *μ*_indel_ < *μ*_bs_ < mtDNA *μ*_bs_ < mtDNA *μ*_indel_. While the small indel rate is lower than the base substitution rate in the nuclear genome, the inverse is true for the mitochondrial genome. A higher indel rate in the mtDNA is largely due to a higher incidence of homopolymeric runs and a greater AT-skew in this genome. In addition, nuclear copy-number changes (gene duplications and deletions) represent a major component of the genetic variation arising due to spontaneous mutation, with rates of origin on the order of 10^−5^ per gene per generation (Konrad et al. 2018).

Our spontaneous nuclear base substitution rate for *C. elegans* of 1.84 × 10^−9^ substitutions/site/generation is similar to two previous estimates for the species using highthroughput sequencing of MA lines (Denver et al. 2009, 2012) but substantially lower than the first estimate which was based on Sanger sequencing (9.1 × 10^−9^; Denver et al. 2004). Additionally, our spontaneous base substitution rate is similar to estimates for the congeneric species *C. briggsae* (average 1.33 × 10^−9^; Denver et al. 2012) and another nematode species, *Pristionchus pacificus* (2.0 × 10^−9^; Weller et al. 2014). The divergence times for *C. elegans*-*C. briggsae* and *Pristionchus*-*Caenorhabditis* are estimated at 80–120 mya (Hillier et al. 2007) and 280-430 mya (Dieterich et al. 2008), respectively. Despite the uncertainty in divergence times based on the molecular clock, the mutation rates of these nematodes under experimental conditions are remarkably similar given the considerable evolutionary time since their divergence, and suggesting that the mutation rates are under stabilizing selection. The base substitution rate in these nematodes is lower relative to other invertebrates for which similar information exists. For example, the base substitution rate in the cladoceran *Daphnia pulex* (Flynn et al. 2017) is roughly twice as high as in nematodes, whereas *D. melanogaster* has an approximately three-fold higher rate than *Caenorhabditis* (Huang et al. 2016; Sharp and Agrawal 2016; Assaf et al. 2017). Furthermore, the spontaneous mitochondrial base substitution rate for the very same *C. elegans* MA lines (Konrad et al. 2017) is 24-fold higher than the nuclear base substitution rate generated from this study.

Spontaneous small indel rates are observed to be considerably lower than base substitution rates for a wide range of surveyed genomes (reviewed in Katju and Bergthorsson 2019). Our spontaneous small indel rate of 6.84 × 10^−10^ changes/site/generation is approximately one-third of the base substitution rate in the *C. elegans* nuclear genome. However, comparing the indel rates with other taxa can be problematic because of the great variation in estimates of indel rates within taxa. For example, indel rate estimates within *D. melanogaster* differ by four-fold whereas the base substitution rates vary less than two-fold (reviewed in Katju and Bergthorsson 2019). Furthermore, many whole-genome sequencing (WGS) studies of MA lines do not report indel rates. However, the small indel rate for *C. elegans* from this study falls within the range reported from MA studies in a few metazoans (0.31 × 10^−9^ to 1.37 × 10^−9^; Katju and Bergthorsson 2019). Our genome-wide estimate of the small indel rate is considerably lower, namely less than 6%, of the originally reported rate for *C. elegans* (Denver et al. 2004). In another notable departure from previous results which found that insertions outnumbered deletions in the *C. elegans* genome (Denver et al. 2004), we find a strong deletion-bias wherein deletions exceed insertions by three-fold. This is consistent with an almost universal deletion-bias observed in MA experiments (reviewed in Katju and Bergthorsson 2019) as well as in comparative analyses of sequenced genomes (Kuo and Ochman 2009). The vast majority of indels occur in homopolymeric runs, and their frequency increases as a function of the length of the run. However, in contrast to A/T runs, short G/C runs appear to have an insertion-bias although long G/C runs have a deletion-bias. Moreover, the indel rates are higher in G/C runs relative to their A/T counterparts. The differences in the mutational properties of low complexity repeats such as homopolymeric runs is likely to play a role in the evolution of their frequency and length distribution in the genome.

The varying population size design of our spontaneous MA experiment allowed us to investigate the influence of increasing selection efficacy on the evolutionary dynamics and persistence of newly occurring nuclear SNP and small indel mutations. Notably, there was no correlation between the frequency of base substitutions, nonsynonymous substitutions, or small indels with population size. This is interesting in light of significant negative correlations observed in this very set of MA lines between population size and (i) nonsynonymous mitochondrial mutations (Konrad et al. 2017), and (ii) many aspects of gene copy-number changes (Konrad et al. 2018). For example, gene deletions accumulated at a higher rate in the *N* = 1 populations than in the larger populations (Konrad et al. 2018). Similarly, both duplications of highly expressed genes, and those that strongly increased the transcript levels of duplicated genes also accumulated more rapidly in the *N* = 1 than in the *N* = 10 or *N* = 100 populations (Konrad et al. 2018). This suggests that both mitochondrial mutations and gene copy-number changes are under more stringent purifying selection than nuclear base substitutions or small indels.

The predominance of transitions over transversions is commonly observed in molecular evolution studies (Vogel and Röhrborn 1966; Fitch 1967; Wakeley 1996). The key mechanisms contributing to this transition bias are held to be (i) selection against transversions which are more likely to cause missense mutations than transitions, and (ii) mutational bias due to the structural similarities among purines and pyrimidines respectively (Stoltzfus and Norris 2016). We did not observe a genome-wide mutational bias towards transitions in our *C. elegans* MA lines, a pattern that has also been noted by others (Denver et al. 2009, 2012). Without any base substitutional bias, transversions are expected to be twice as frequent as transitions and the frequency of transitions and transversions in our study is not significantly different from this expectation. However, in exons where a transition/transversion bias is most likely to have consequences for fitness, we do in fact observe a transition bias. The number of transitions and transversions are roughly equal in exons, which means that transitions are twice as frequent as expected if there was no bias. The near universal base substitution bias towards A/T nucleotides is also observed in our results as G/C → A/T substitutions are 3.75-fold more likely than mutations in the opposite direction. This base substitution bias predicts an equilibrium base composition of 26% G/C, which is lower than either total G/C content of the *C. elegans* genome (36%) or the G/C content of intergenic DNA and introns (33%). Assuming that the mutational biases under experimental condition are the same as the prevailing mutational biases in the wild, the departure of the observed G+C-content from the expected suggests that other mechanisms than the biases of spontaneous mutations are influencing the base composition of the *C. elegans* nuclear genome. Higher G+C-content than expected by mutation pressure alone seems to be the rule in genome evolution, and it is usually presumed that natural selection for higher G+C-content and/or biased gene conversion are responsible. However, this departure from equilibrium G+C-content also has the effect of increasing the mutation rate (Krasovec et al. 2017).

Furthermore, there are interesting context-dependent patterns in the frequency of substitutions. In particular, a 5’–T and 3’–A have a strong positive effect on the A/T → T/A substitution rate, especially at the boundaries of A or T homopolymeric runs. Similar observations have been made in mismatch-repair deficient lines of *C. elegans* (Meier et al. 2018). The combination of this strong context-dependence of base substitutions and the genomic distribution of A and T homopolymeric runs explains three other observations about the base substitution patterns in our MA lines. Introns and intergenic regions have significantly higher mutation rates than exons in our study. It is usually assumed that differences in substitution rates between introns and exons are due to selection rather than intrinsic differences in mutation rates. However, lower mutation rates in coding sequences relative to non-coding ones have been observed in other MA experiments and were ascribed to transcription-coupled repair (TCR) and differential efficiency of mismatch repair (MMR) between coding and non-coding DNA (Krasovec et al. 2017). Additionally, a recent study of somatic mutation rates in humans concluded that introns have higher mutation rates than exons due in part to greater efficiency of mismatch repair in exons (Frigola et al. 2017). The data presented here suggest that the difference in mutation rates between introns and exons in *C. elegans* is caused by strongly context-dependent A/T → T/A substitution mutations. These mutations, which are particularly frequent at the boundaries of A and T homopolymeric runs, are in turn more common in introns and intergenic regions and less prevalent in exons. Indeed, if we exclude A/T → T/A mutations from our analysis, the difference in mutation rates between exons and introns disappears. Hence, the higher mutation rates in introns and intergenic regions compared to exons in *C. elegans* is due to a higher prevalence of mutagenic motifs in introns and intergenic regions.

Nucleotide polymorphisms in natural populations are correlated with recombination rates (Begun and Aquadro 1992; Cutter and Choi 2010; McGaugh et al. 2012). These correlations are usually attributed to the combination of natural selection and genetic linkage where genetic hitchhiking or background selection on linked sites depresses genetic variation in regions of low recombination. However, mutation rates are also positively correlated with recombination rates in several well-studied systems such as humans, *Arabidopsis*, honey bees and *C. elegans* (Arbeithuber et al. 2015; Francioli et al. 2015; Yang et al. 2015; Konrad et al. 2018; Smith et al. 2018). The *C. elegans* chromosomes can be divided into three regions with respect to recombination frequency (Rockman and Kruglyak 2009). The most central regions of the chromosomes, the cores, have low recombination frequency, the arms have high recombination frequency, and the tips have low recombination frequency. Our previous study of spontaneous gene copy-number changes in these *C. elegans* MA lines found that duplication and deletion breakpoints were more frequent in arms and tips than in the cores (Konrad et al. 2018). In this study, the distribution of base substitutions and indels follow the same pattern, with significantly lower mutation rates in the cores relative to the arms and tips. Our comparison of the base substitution spectrum in cores *vs.* arms and tips revealed that A/T → T/A mutations are disproportionately more common in the arms and tips than in the cores. Even when A/T → T/A mutations are excluded from the analysis, there is still a difference in substitution rates between recombination domains. However, just as with the difference in mutation rates between exons, introns and intergenic regions, the difference in mutation rates between cores *vs.* arms and tips is also a function of the frequency of A/T homopolymeric runs.

Experiments in several organisms have suggested that frequent transcription can render the transcribed DNA more vulnerable to mutations (Klapacz and Bhagwat 2002; Hudson et al. 2003; Kim and Jinks-Robertson 2012). For such an effect to influence the mutation rates in multicellular animals, germline transcribed genes could hypothetically have higher mutation rates than genes that are only expressed in the somatic tissues. Our results initially suggested that germline expressed genes may have higher substitution rates than non-germline expressed genes. However, this effect was only detected in germline transcribed genes located in the chromosomal arms, and not in the cores. Upon further analysis, we found that the association between germline transcription and the base substitution was due to context-dependent A/T → T/A substitutions in the introns of germline transcribed genes. Hence, the higher mutation rates of germline expressed genes in our MA lines was not due to a general increase in the substitution rate and it did not extend to exons of these genes.

This study contains the largest set of mutations for a spontaneous MA experiment employing the *C. elegans* N2 wild-type strain. The analysis of base substitutions in our MA lines confirmed some previous results regarding the mutation rates, and mutational biases. Other results add context to previous observations. For example, the lack of transition bias is primarily due to high transversion rates, specifically A/T → T/A, in introns and intergenic regions and does not extend to exons. The analysis also illustrates that correlations between recombination frequency, genomic location and transcription with mutation rate can arise from the nonrandom distribution of mutagenic motifs. The efficacy of natural selection versus genetic drift depends on the effective population size. These MA experiments utilized different population sizes to reveal the effects of different efficacy of selection on the accumulation of mutations. Previous phenotypic analyses of these MA lines for two fitness-related traits indicated that (i) the *N* = 10 and *N* =100 populations did not suffer significant decline in fitness due to deleterious mutations, and (ii) most of the decline in fitness in the *N* = 1 populations was due to mutations of large effects (Katju et al. 2015, 2018). Alternatively, the observed decline in fitness traits could be due to a large number of mutations with small fitness effects. The lack of a correlation between nuclear base substitution rates and population sizes is consistent with the previous results that a small number of mutations are responsible for the fitness decline in the *N* = 1 lines. Finally, we note that a negative correlation was indeed found between population size and the accumulation of mitochondrial mutations, gene deletion rates and transcript abundance of duplicated genes in these experiments. The differences between the results for mitochondrial mutations and gene copy-number changes on the one hand, and nuclear base substitutions and small indels, on the other, are consistent with the view that the former have, on average, more detrimental effects on fitness.

## METHODS

### Mutation accumulation experiment

As a self-fertilizing nematode with a generation time of 3.5 days at 20 °C, and the ability to survive long-term cryogenic storage, *C. elegans* is an ideal organism for MA studies. The spontaneous MA experiment was initiated with a single wild-type Bristol (N2) hermaphrodite originally isolated as a virgin L4 larva. The F1 hermaphrodite descendants of this single worm were further inbred by self-fertilization before establishing 35 MA lines and cryogenically preserving thousands of excess animals at −86**°**C for use as ancestral controls. 20 of these 35 lines were established with a single worm and propagated at *N* = 1 individual per generation. Ten lines were initiated with ten randomly chosen L4 hermaphrodite larvae and subsequently bottlenecked each generation at *N* = 10. Five lines were initiated and subsequently maintained each generation with 100 randomly chosen L4 hermaphrodite larvae (*N* = 100). A new generation was established every four days. The *N* = 1, 10 and 100 population size treatments correspond to effective population sizes (*N_e_*) of 1, 5, and 50, respectively (Katju et al. 2015, 2018). The worms were cultured using standard techniques with maintenance at 20**°**C on NGM agar in (i) 60×15 mm Petri dishes seeded with 250 *μl* suspension of *E. coli* strain OP50 in YT media (*N* = 1 and *N* = 10 lines) or (ii) 90×15 mm Petri dishes seeded with 750 *μl* suspension of E. coli strain OP50 in YT media (*N* = 100 lines). Stocks of the MA lines were cryogenically preserved at −86**°**C every 50 generations. The experiment was terminated following 409 MA generations because the *N* = 1 lines displayed a highly significant fitness decline. Three lines were already extinct due to the accumulation of a significant mutation load and five additional lines were on the verge of extinction (displaying great difficulty in generation to generation propagation).

### DNA preparation and sequencing

Following the completion of the MA phase, a total of 86 worms were prepared for DNA whole genome sequencing: one worm from every population of size *N* = 1, four individuals from every population of size *N* = 10, five individuals from every population of size *N* = 100, and one individual from the ancestral strain used to set up the MA experiment Each of the 86 individuals were allowed to go through several self-fertilization and reproductive cycles to generate enough offspring necessary for genomic DNA extraction. The preparation for sequencing followed the methodology previously described (Konrad et al. 2017, 2018). Genomic DNA was isolated with the PureGene Genomic DNA Tissue Kit (QIAGEN no. 158622) and a supplemental nematode protocol. The quality and concentration of the gDNA were checked on 1% agarose gels via electrophoresis, BR Qubit assay (Invitrogen), and a Nanodrop spectrophotometer (Thermo Fisher). Target fragment lengths of 200-400bp were prepared via sonication of 2*μg* of each DNA sample in 85*μl* TE buffer, end-repaired (NEBNext end repair module (New England BioLabs)) and purified (Agencourt AMPure XP beads (Beckman Coulter Genomics)). Beads used during the purification were not removed until after adapter ligation as has been described previously (Thompson et al. 2013). Custom pre-annealed Illumina adapters were ligated to the fragments and 3’ adenine overhangs were added (AmpliTaq DNA Polymerase Kit, Life Technologies). Kapa Hifi DNA Polymerase (Kapa Biosystems) with Illumina’s paired end genomic DNA primers containing 8 bp barcodes was used for PCR amplification. PCR products were size fractionated on 6% PAGE gels and 300-400bp fractions were selected for excision. The fragments were gel extracted via diffusion at 65°C and gel filtrated (NanoSep, Pall Life Sciences). A final purification step was performed using Agencourt AMPure beads. The final DNA quality and quantity were evaluated using the Agilent HS Bioanalyzer and HS Qbit assays. The multiplexed DNA libraries were sequenced on Illumina HiSeq sequencers with default quality filters at the Northwest Genomics Center (University of Washington).

### Sequence alignment and identification of putative variants

The demultiplexed raw reads stored as individual fastq files for each genome were aligned to the reference N2 genome (version WS247; www.wormbase.org; Harris et al. 2010) via the Burrows-Wheeler Aligner (BWA Version 0.5.9) (Li and Durbin 2009) and via Phaster (Green lab) and prepared for analysis as previously described (Konrad et al. 2018).

Seventeen lines of size *N* = 1 were included in the final analysis (1A-1H, 1K, and 1M-1T). The alignment files were used to identify all putative base substitution and indels within the 82 individual descendants relative to the ancestral genome. Putative substitutions and indels were identified separately for the Phaster and BWA alignments using Platypus (Rimmer et al. 2014), Freebayes (Garrison and Marth 2012), and a pipeline consisting of mpileup (Li et al. 2009), bcftools (Li 2010), vcfutils (Danecek et al. 2011) and custom filters written in Perl. Indel calls were based primarily on Phaster alignments, but were verified in the BWA alignments. Indelminer (Ratan et al. 2015) was used as an additional approach to call indels with the ancestral line as a direct reference. A minimum root-mean-square mapping quality of 30 was required for SNPs to be retained, while a mapping quality of 40 was required for indels. SNPs were required to have a minimum support of three quality reads, while indels were required to be covered by a minimum of five quality reads. Variants that occurred even with low quality or coverage in the ancestral line were removed from the analysis. Only variants supported by at least 80% of the high-quality reads at its position were retained in the dataset. Each variant had to be confirmed by at least two of the variant callers in order to be considered for further analysis.

### Binomial Probability Verification

Every variant was independently verified by calculating a binomial probability for it, given the number of variant calls at the same location in the genome across all other genomes sequenced. For each putative variant position, the number of read calling the same variant were summed. For each putative variant position, the number of reads across all lines calling the variant were summed and divided by the total number of reads at the variant position. We used this as the probability of any given read calling the variant by chance (*P*). For each putative mutation, we counted the number of reads within every individual line which called the variants (*K*), and the total number of reads at the position in that line (*N*). We then calculated the p-value for the variant (*var*) in that line (*i*):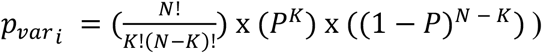. The probabilities across all lines where sorted from most significant to least significant, and a Holm-Bonferroni correction was applied to determine if the variants called by the previous pipeline met the critical p-value threshold.

### Independent validation of SNP and Small Indel Variants

All substitutions and indels identified in the exons of the *N* = 1 lines were checked against the RNA-sequencing data set previously described in Konrad et al. (2018). The RNA-Seq reads were realigned using STAR in order to allow for indel-aware alignment of these reads (Dobin et al. 2013). Verification of all variants was done via computational analysis of the CIGAR scores in the BAM files, and finalized manually using the Integrative Genomics Viewer (Thorvaldsdóttir et al. 2013). Of the 199 substitutions detected in the exons, 195 were verified by RNA-Seq data. The four variants that could not be validated by RNA-Seq were associated with line 1T which went extinct at MA generation 309 (Katju et al. 2015, 2018). RNA for line 1T was extracted from an earlier stock cryopreserved at MA generation 305. 35 indels were detected in exons that were also covered by the RNA-Seq data. All of these indels were verified in the RNA-Seq data.

In addition, we randomly selected 46 SNP and small indel variants identified by whole-genome sequencing in the introns and intergenic regions of the 17 *N* =1 MA lines for independent confirmation via PCR and Sanger sequencing. Primers were designed to amplify regions containing candidate mutations. The locus of interest was sequenced in the candidate MA line as well as the ancestral control. PCR products were purified using a silica membrane protocol and Sanger sequenced by Eton Biosciences Inc. Sequences were mapped to the reference genome using BLAST and alignments were inspected to verify either the ancestral sequence or new variant. Chromatograms were examined to ensure sequence quality. 44 of the 46 variants were independently validated using this approach. Two mutations in MA line 1T could not be verified. Both these mutations were initially detected within segmental duplications. This line demonstrated evidence of chromothripsis and went extinct prior to the termination of the Ma experiment, which may have been a complicating factor (Konrad et al. 2018).

### Annotation, Characterization, and Mutation Rate Calculations for SNPs and Indels

All variants were annotated based on the GFF file available for the N2 reference genome of *C. elegans* (version WS247; www.wormbase.org; Harris et al. 2010) using a custom script. Mutations were assigned to exons, introns, and intergenic regions (if the mutation fell outside a protein coding gene), and to chromosomal arms, cores, and tips based on boundaries predicted by Rockman and Kruglyak (2009). The mutation rate 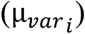 was estimated individually for each population as variants (or sum of variant frequencies) per base per generation 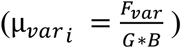, where *F_var_* refers to the number (or sum of frequencies) of single nucleotide polymorphisms or indels within the line, G refers to the number of generations through which the line was propagated, and B_*total*_ refers to the total number of bases in the genome that meet the same thresholds required for variant identification relative to the N2 reference genome (version WS247). B_*total*_ was individually calculated for each genome by counting the number of positions within the sequenced genome that met the same quality thresholds as those required for a variant to be called. For populations of size *N* > 1, the sum of frequencies of variants was calculated from the proportion of individuals sequenced for each population that carried each of the variants of interest. B_*total*_ in populations of size *N* > 1 was averaged across the genomes of the individuals sequenced for that population. Mutation rates for each of the population sizes were calculated by averaging the population-specific mutation rates within each population size treatment: 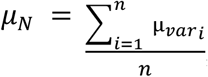, where *var_i_* refers to the population specific mutation rate, and n refers to the total number of populations of a given population size (*N*) (17, 10, and 5 for populations of size *N* = 1, 10, and 100, respectively). The number of generations through which each population was propagated differed between the lines of size *N* = 1 (Supplementary Table 2), as some populations became too sick to be propagated any further, or went extinct.

Every mutation was initially assigned to one of three intra-chromosomal regions classified by their recombination rates as described in Rockman and Kruglyak (2009): cores, arms, and tips. The expected distribution of variants across these regions was estimated based on the proportion of the genome falling within each category. Every protein coding gene was categorized as either a germline or non-germline expressed gene based on the data of Wang et al. (2009). Germline mutation rates were calculated by summing the number of mutations within each line that fell onto any of the germline genes and dividing that by the total number of high-quality bases within germline genes. Mutation rates for non-germline genes were calculated in the same fashion.

We calculated the median amino acid radicality for the pool of amino acid replacement substitutions by first calculating a radicality score for each amino acid change. For this, we used the six biochemical classification schemes described in Sharbrough et al. (2018) to determine how radical any given amino acid change is. For instance, if a pair of amino acids is assigned into the same class for all six schemes, the amino acid substitution is assigned a score of 0. If only three out of the six schemes assign the amino acids into the same category, the substitution will have a score of 0.5, and if no scheme classifies the amino acids the same, the substitution will have a radicality of 1. Before the mean of the radicality scores for each substitution within a line was calculated, we normalized each score by the frequency of the variant within its population.

Normalization of mutation spectra and category specific mutation rates (arms, cores, tips, exons, introns, etc.) were calculated by dividing the raw variant counts or frequencies for each category by the number of bases in the genome belonging to each category and which met the same quality thresholds as those required for variant calling.

Sequence complexity was calculated as previously described (Morgulis et al. 2006). Briefly, given a sequence (*a*) of length *n* and 64 possible triplets of {A, C, G, T}, the occurrence of each possible triplet (*t*) was counted across the sequence and yields *c_t_(a)*. The total number of overlapping triplets occurring in any sequence (*l*) equals *n*-2. Sequence complexity (*S(a)*) was then calculated as:

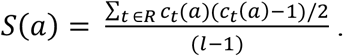

All statistical tests were performed in R (R Core Development Team 2014).

## DATA ACCESS

Sequence data from the MA experiment in this has been deposited under NCBI BioProject PRJNA448413.

## ACKNOWLEDGEMENTS

We thank Lucille Packard for assistance in the creation of the MA lines, and Philip Green from the University of Washington for providing the program Phaster. This research was supported by National Science Foundation Grant MCB-1330245 to V.K. U.B. and V.K. were additionally supported by start-up funds from the Department of Veterinary Integrative Biosciences, College of Veterinary Medicine and Biomedical Sciences at Texas A&M University.

